# mRNA adenosine methylase (MTA) deposits m^6^A on pri-miRNAs to modulate miRNA biogenesis in *Arabidopsis thaliana*

**DOI:** 10.1101/557900

**Authors:** Susheel Sagar Bhat, Dawid Bielewicz, Natalia Grzelak, Tomasz Gulanicz, Zsuzsanna Bodi, Lukasz Szewc, Mateusz Bajczyk, Jakub Dolata, Dariusz J. Smolinski, Rupert G. Fray, Artur Jarmolowski, Zofia Szweykowska-Kulinska

## Abstract

m^6^A, one of the most abundant mRNA modifications, has been associated with various metabolic processes in plants. Here we show that m^6^A also plays a role in miRNA biogenesis in *Arabidopsis thaliana.* Significant reductions in plant m^6^A/MTA levels results in lower accumulation of miRNAs whereas pri-miRNA levels tend to be higher in such plants. m^6^A-IP Seq and MTA-GFP RIP were used to show that many pri-miRNAs are m^6^A methylated and are bound by MTA, further demonstrating that pri-miRNAs can also be substrates for m^6^A methylation by MTA. We report that MTA interacts with RNA Pol II, supporting the assumption that m^6^A methylation is a co-transcriptional process, and also identify TGH, a known miRNA biogenesis related protein, as a novel protein that interacts with MTA. Finally, reduced levels of miR393b may partially explain the strong auxin insensitivity seen in *Arabidopsis* plants with reduced m^6^A levels.

## Introduction

*N*^6^-methyladenosine (m^6^A), one of the most abundant mRNA modifications in eukaryotic cells can regulate eukaryote gene expression at multiple post- and co-transcriptional levels. m^6^A methylation in animal mRNAs is associated with several biological processes, ranging from cancer^1^, viral infections^2,3^ to cell development^4,5^ with the underpinning mechanisms including m^6^A regulated pre-mRNA splicing patterns, mRNA export, mRNA stability and changes in translational efficiency^6^. A group of proteins that collectively form the RNA methylation “writer” complex have been characterized and are well conserved between plants and animals. The mammalian m^6^A methyltranserase complex consists of Methyltransferase Like 3 (METTL3)^7^, Methyltransferase Like 14 (METTL14)^8^, Wilms’ Tumour1-Associating Protein (WTAP)^9^, VIRMA (KIAA1429)^10^, RNA-binding motif protein 15 (RBM15)^11^ and Zinc Finger CCCH-Type Containing 13 (ZC3H13)^12,13^. While METTL3 has been identified as the catalytic protein in this complex^7^, auxiliary proteins provide specificity and/or help with proper localization of the complex^6^. The m^6^A mark can be recognized by various “readers”, the best characterized of which belong to the YT521-B homology (YTH) domain family^14-17^. The modification can also be removed from transcripts by “erasers”, which in humans include fat mass and obesity-associated protein (FTO)^18^ and α-ketoglutarate-dependent dioxygenase alkB homolog 5 (ALKBH5)^19^.

In *Arabidopsis thaliana,* the presence of m^6^A was first reported in 2008 and was shown to be dependent upon the activity of mRNA adenosine methylase (MTA) [homolog of human METTL3], the catalytic component of *Arabidopsis* m^6^A methyltransferase complex^20^. FKBP12 interacting protein 37 kDa (FIP37, homolog of WTAP) was the first identified methyltransferase interacting complex member in eukaryotes^20^. Later, FIP37 was further characterised and shown to be required for m^6^A formation in mRNA^21^. Other protein components of m^6^A methyltransferase complex that have been identified include MTB (Methytransferase B, homolog of METTL14), VIR (Virilizer, homolog of VIRMA) and *At*HAKAI (homolog of HAKAI)^22^. In *Arabidopsis,* Evolutionarily Conserved C-Terminal Region (ECT), are the orthologues of the mammalian YTH family of proteins and are the only readers identified so far^23-25^. ALKBH10B and ALKBH9B represent the characterized *Arabidopsis* m^6^A erasers ^26,27^.

Null mutants of MTA or any other members of the core writers complex i.e. MTB, FIP37 and VIR are embryo lethal ^20-22^, indicating an essential function for this modification. However, hypomorphic knockdown mutants of these writers have been obtained and typically show an 80-90% reduction in m^6^A levels^21,22,28^. In addition, reader and eraser knockouts are viable^25-27^. Studies utilizing these various mutant plant lines indicate a role of this modification in embryogenesis, proper plant development (trichome morphology, meristem maintenance, vascular development), flowering time and flower morphology, and pathogen response^20,22,26-29^.

Several aspects of plant development and metabolism are controlled by micro RNAs (miRNAs), and the complex phenotypes of low methylation plants have aspects reminiscent of miRNA biogenesis pathway mutants (review^30^) and might be partially explained by m^6^A controlled miRNA biogenesis. miRNAs are small endogenous non-coding RNAs that are ∼21nt in length and are important players in regulating cellular metabolism. miRNAs are derived from hairpin loop containing pri-miRNAs that are transcribed by RNA Polymerase II, and further processed by RNAse III type enzymes associated and assisted by other proteins. In animals, where m^6^A is important for biogenesis of many microRNAs, m^6^A plays the role of a mark, identified by a reader protein (Heterogeneous Nuclear Ribonucleoprotein A2/B1; HNRNPA2B1) that facilitates the recruitment of downstream enzymes (RNAse III and associates) to pri-miRNAs thus facilitating their processing to miRNAs. Accordingly, it was shown that depletion of METTL3 leads to decreased accumulation of miRNAs and to an over accumulation of pri-miRNAs due to their impaired processing^31,32^. While most of the findings related to m^6^A in plants concern mRNAs, its role in miRNA biogenesis in plants remains unknown.

In this study, we provide evidence that biogenesis of at least 25% of Arabidopsis miRNAs is affected by the presence of m^6^A mark. We show that plant pri-miRNAs are m^6^A methylated by MTA, and deficiency of MTA (and thus m^6^A) leads to accumulation of pri-miRNAs accompanied by lower miRNA levels. We also show that MTA interacts with RNA Pol II and Tough (TGH) suggesting that MTA acts at early stages of miRNA biogenesis. We also suggest that the impaired auxin response in plants with MTA deficiency is caused, at least partly, by decreased miR393b levels.

## Results

### Lack of m^6^A results in impaired miRNA biogenesis

In order to assess possible effects of m^6^A levels on miRNA biogenesis, we performed small RNA (sRNA) sequencing on rosette leaves from 4 weeks old plants from both wild type (WT) Col-0 and a mutant line with severely reduced m^6^A levels. The reduced *mta* line contains MTA cDNA under the ABI3 promoter in a homozygous MTA T-DNA insertion mutant line. The ABI3 promoter drives strong embryo expression of MTA, enabling the plant embryo lethality to be bypassed. However, this promoter drives a very low level of expression post germination, giving rise to plants with 80-90% less methylation compared to their WT counterparts^28^. Our sequencing data showed a decrease in overall abundance of mature miRNAs (**Fig. S1**) and out of these we were able to identify 60 differentially expressed miRNAs that had high confidence score (probability ≥ 0.9 or false-discovery rate FDR = 0.1). 51 out of these 60 miRNAs were found to be downregulated while only 9 miRNAs were upregulated (**Fig. 1a**). This downregulation of miRNAs in *mta* plants was also confirmed by quantitative real time PCR (RT-qPCR) for 6 randomly selected miRNAs (miR159b, miR169a, miR319b, miR396b, and mir399a and miR850)(**Fig. 1b**). Following from these results we went on to investigate the levels of primary-miRNAs (pri-miRNAs) in the low methylation plants by using the mirEX^2^ platform (http://www.combio.pl/mirex2), developed by our group^33,34^. This platform utilizes a repository of 298 primers, specifically for pri-miRNA quantification by qRT-PCR. Out of 298 pri-miRNAs tested, 68 pri-miRNAs were at undetectable levels in our samples leaving 230 for further analysis. Results from RT-qPCR revealed that 85 pri-miRNAs had statistically significant (p-value = <0.05, n = 3) changes in their accumulation (WT vs *mta*); 56 of these (∼66%) were found to be upregulated while 29 were downregulated (**Fig. 1c**). On comparison of the RT-qPCR results with sRNA sequencing results we were able to identify 20 cognate pri-miRNA/miRNA pairs that showed higher accumulation of pri-miRNAs and lower miRNA levels (**Fig. 1d)**. These results point towards an impaired processing of pri-miRNAs in the absence of MTA.

**Figure 1.**
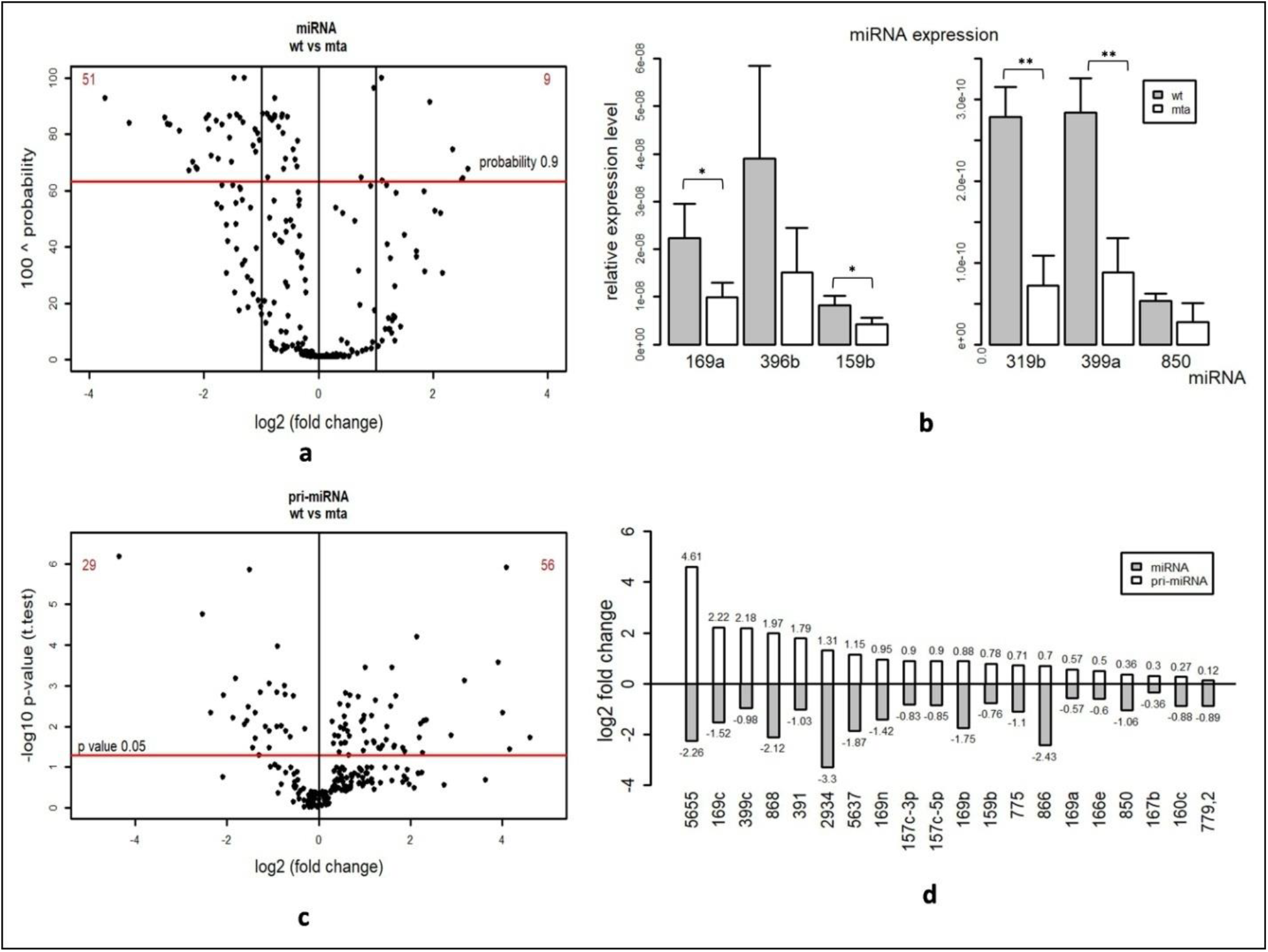
miRNA biogenesis is impaired in *mta* mutants. **a)** Small RNA sequencing analysis of miRNAs in *mta* and WT plants. Each black dot represents one miRNA. Red horizontal bar represents the threshold (probability 0.9). **b)** Relative abundance (as determined by TaqMan RT-qPCR) of miRNAs identified with altered abundance by sRNA Seq in WT vs *mta* mutant. * = p-value < 0.05, ** = p-value < 0.005 and error bars represent standard deviation (n=3). **c)** Levels of 260 pri-miRNAs as determined by RT-qPCR with separation of statistically significant pri-miRNAs (above red horizontal bar). Each dot represents one pri-miRNA and red horizontal bar represents p-value threshold (p-value 0.05). **d)** A set of cognate pairs of pri-miRNAs/miRNAs (selected from panel **a** and **b**) where pri-miRNA levels are upregulated and miRNA levels are downregulted.

### *Arabidopsis* pri-miRNAs are m^6^A methylated by MTA

The increased level of pri-miRNAs in the low methylation plants raised the possibility that the presence of m^6^A is necessary for proper processing of some pri-miRNAs. Therefore, we set out to test the methylation status of the pri-miRNAs using m^6^A-RNA immunoprecipitation followed by sequencing (m^6^A-IP Seq). This method is not an equivalent of the commonly used MeRIPSeq^35^, as we avoided the fragmentation of polyA RNA. We validated this protocol using spiked-in methylated and non-methylated controls (**Fig. S2**). From the m^6^A-IP Seq data we selected miRNA genes (*MIR*) that were independent transcriptional units (not in introns of protein coding regions) and were either absent in *mta* samples or had significantly reduced abundance relative to WT Col-0. Using this approach, we identified transcripts of 11 *MIR* genes that were enriched more than 1.5 folds in WT Col-0 vs *mta*, indicating that they are m^6^A methylated in WT plants (hence can be immunoprecipitated with m^6^A antibody) and in the absence of MTA, lack this methylation (**Fig 2a, S3**). We further checked the robustness of our data by comparing it to previously published data by Shen *et. al*.^21^ and Anderson *et. al.*^36^. In our data set, we were able to identify 14,870 genes whose transcripts (15,562) are depleted in *mta* mutant indicating that these gene transcripts are methylated. Upon comparison we found an overlap of 2,868 and 3,100 genes between our data and data from Shen *et. al.* and Anderson *et. al.,* respectively. 930 genes were common among all three datasets (**Fig. 2b**).

**Figure 2.**
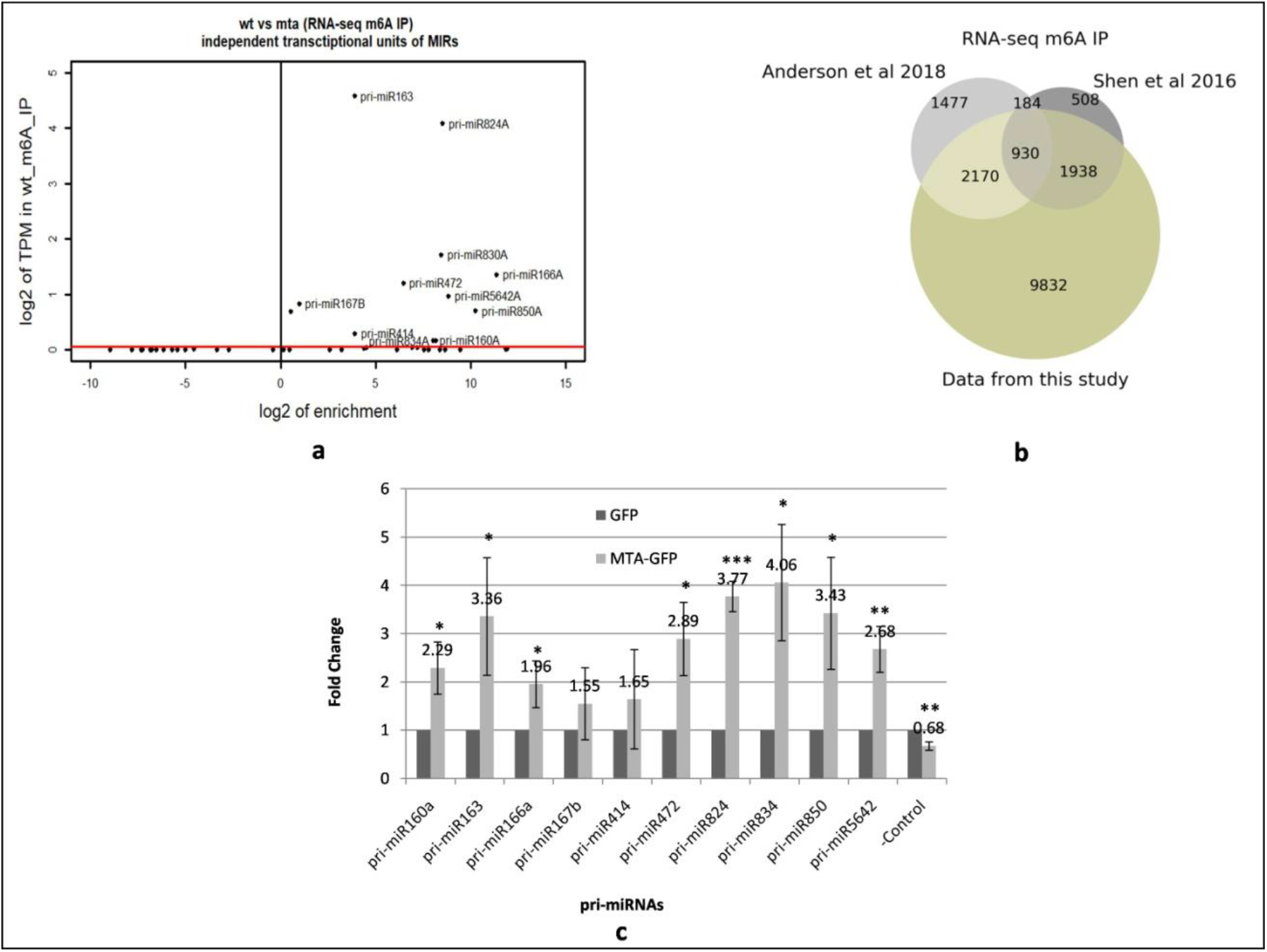
pri-miRNAs are m^6^A methylated by MTA. **a)** *MIR* gene precursors identified by m^6^A IP Seq. Red bar represents the threshold of TPM > 0.05. **b)** Comparison of our m^6^A-IP Seq data with data from m^6^A-IP obtained from other *Arabidopsis* mutants of m^6^A writer proteins as described by Shen *et. al.* and Anderson *et. al*^21,36^. Numbers represent the number of genes identified in each experiment. **c)** Levels of pri-miRNAs in MTA-GFP samples after GFP trap-based RIP RT-qPCR are presented as fold changes as compared to GFP control RIP RT-qPCR samples. * = p-value < 0.05, ** = p-value < 0.005, *** = p-value < 0.001. Error bars represent standard deviation of 3 biological replicates. – Control = *AT2G40000* gene.

As an orthogonal approach to verify that these pri-miRNAs were bona fide targets for MTA directed methylation, we carried out RNA immunoprecipitation (RIP) using an anti-GFP antibody and GFP tagged MTA (35S:*MTA*:GFP) plant lines. RIP was performed on the nuclear fraction and followed by RT-qPCR on oligo(dT) primed cDNA. 18 pri-miRNAs (10 from our m^6^A-IP Seq data and 8 random) were selected for checking by RT-qPCR, of these, all but four showed a statistically significant enrichment (p-value < 0.05) in the MTA-GFP sample (**Fig. 2c, S4**). This indicates that pri-miRNAs are commonly bound by MTA. Taken together, these data suggest that, at least for some pri-miRNAs, MTA binds and methylates the transcript and that this promotes their processing to mature miRNAs.

### MTA interacts with RNA Pol II *in situ*

The accumulation of pri-miRNAs and low abundance of miRNAs in *mta* mutants is reminiscent of various plant miRNA biogenesis protein mutants like Dicer like 1 (DCL1), Hyponastic Leaves 1 (HYL1)^37-39^, Serrate (SE)^40^ and Tough (TGH)^41^. These proteins are involved in early stages of miRNA biogenesis and are needed for efficient processing of pri-miRNAs to miRNAs. These observations led us to believe that MTA might act at early stages of miRNA biogenesis, maybe even co-transcriptionally. The deposition of m^6^A is usually assumed to be co-transcriptional, not least because it can influence other processing events such as poly adenylation site choice or pre-mRNA splicing^42-44^, which are themselves often co-transcriptional. However, a direct association between METTL3 and RNA Pol II has only been shown in mammalian systems when the rate of transcription was artificially slowed^45^. We confirmed colocalization of MTA and RNA Pol II in *Arabidopsis* using immunolocalzation (**Fig. S5**). Furthermore, we used proximity ligation assay (PLA) to confirm *in-vivo* interactions of MTA with RNA Pol II in plants under physiological conditions. Our results show direct interactions between MTA-GFP and RNA Pol II in cell nuclei with MTA-GFP expression but not in GFP control (**Fig 3**). These results were obtained for RNA Pol II phosphorylated at both Serine 5 and Serine 2. Thus, we show that MTA is associated with the RNA Pol II from an early stage of transcription and likely methylates pri-miRNAs (among other RNA Pol II transcripts) co-transcriptionally.

**Figure 3.**
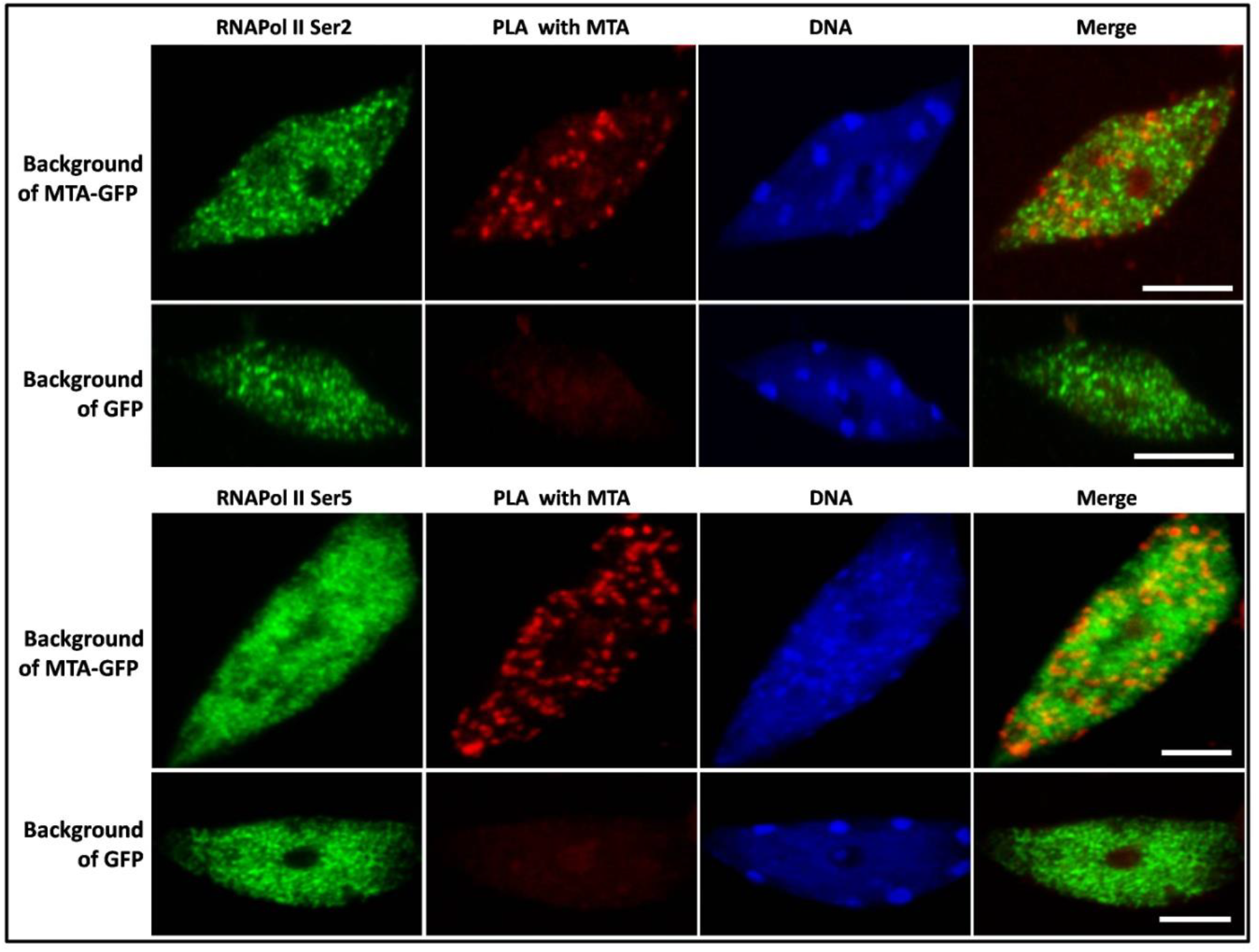
PLA shows interaction between MTA and RNA Pol II phosphorylated at Serine 5 and Serine 2. Positive PLA signals (seen in second column) can be seen only in cells containing MTA-GFP transgene but not in control cells containing GFP transgene control samples. RNA Pol II is represented in green. DNA is stained with HOECHST (blue). Scales bars (white) = 5µm.

### MTA acts at early stages of miRNA biogenesis

While MTA interacting with RNA Pol II is an indication of it methylating RNA Pol II transcripts co-transcriptionally in general, we investigated whether any protein involved in miRNA biogenesis would also interact with MTA and modulate the process via m^6^A methylation. To address this possibility, we performed a screen on possible MTA interactors that are known players in miRNA biogenesis, like HYL1, Cap binding protein 20/80 (CBP20/80), SE and TGH etc., using Yeast Two Hybrid system. In this screen we identified TGH as a positive MTA interactor (**Fig. S6**) and these results were later supported using microscopy to demonstrate co-localisation of GFP-TGH and RFP-MTA in *Arabidopsis* protoplasts (**Fig. 4a**). When TGH and MTA are expressed together, MTA is also present in nuclear speckles that are formed by TGH, indicating a strong association between the two. To confirm this interaction, we performed Forster resonance energy transfer (FRET) analyzed by Fluorescence Lifetime Imaging Microscopy (FLIM). As FRET can occur only when two proteins are within nanometers of each other, FRET-FLIM helps to quantify direct protein interactions. FRET-FLIM analysis further confirmed the close association and interaction of these proteins (**Fig. 4b**). Thus, we have identified TGH as a novel MTA partner. This discovery led us to analyze the small RNA sequencing data from *tgh* mutant that is already available^41^ (accession no.: GSE38600); although this data was obtained from inflorescence we found that aside from the overall downregulation of miRNAs, 23 miRNAs were commonly downregulated in both *mta* and *tgh* mutants. While we did not expect a big overlap between these two data sets owing to different source plant tissue, an overlap of 45% (23 out of 51) is still substantial and indicates that MTA and TGH may play a role together in processing, at least for a set of pri-miRNAs (**Fig. S7**).

**Figure 4.**
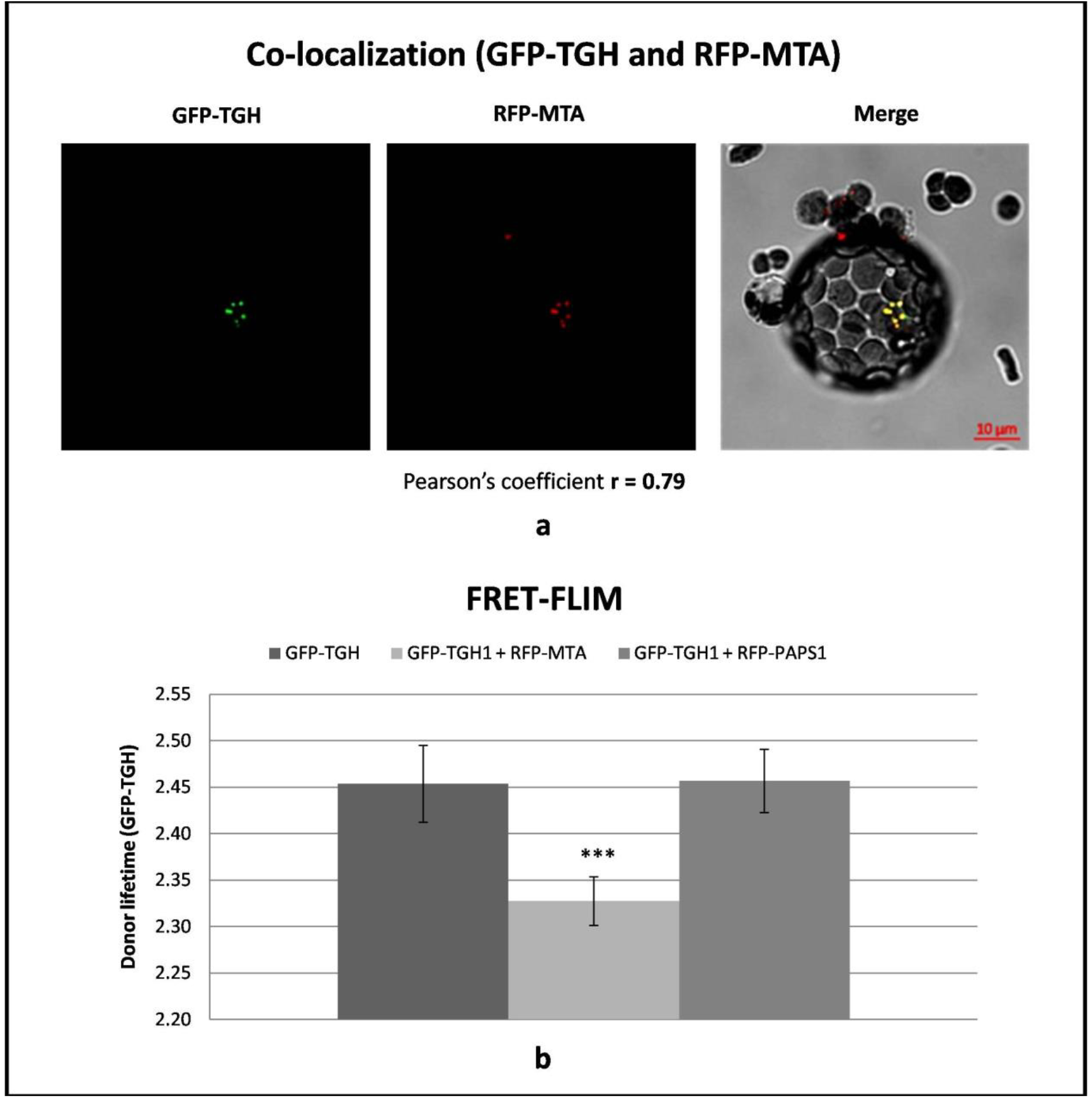
MTA interacts with miRNA biogenesis related protein, TGH. **a)** *Arabidopsis* WT protoplasts co-transfected with GFP-TGH and RFP-MTA (left and middle panel) showing co-localization of both proteins in the nucleus. Right panel presents merged image and scale bar (10µm) is shown in red. **b)** FRET-FLIM analysis in *Arabidopsis* protoplasts co-transfected with GFP-TGH and RFP-MTA or GFP-TGH and RFP-PAPS1. Reduction in donor lifetime (GFP-TGH) was seen with RFP-MTA but not with negative control (RFP-PAPS1); *** = p-value <0.001, n = 9.

### MTA regulates the level of miR393b which is involved in auxin response

Auxin response defects have been reported for hypomorphic mutants of the known plant m^6^A writers^22^, indeed, we found this to be the case for the auxin responsive *DR5pro:GUS* reporter construct when introduced into both WT Col-0 and the *mta* mutant background. Upon induction of 14-day old seedlings with 2,4-dichlorophenoxyacetic acid (2,4-D) *mta* plants showed much less GUS expression (**Fig. 5a**).

**Figure 5.**
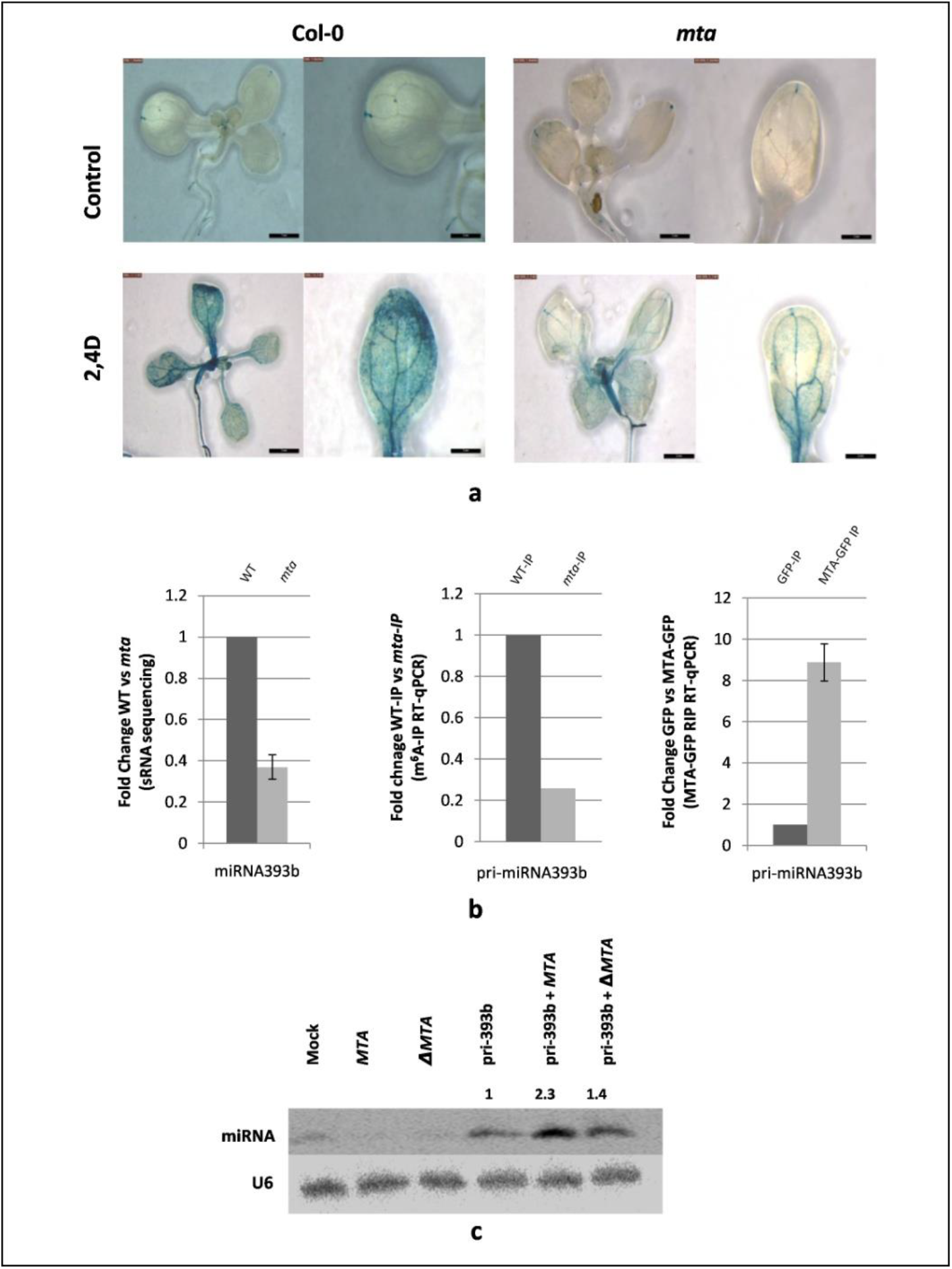
Auxin insensitivity in *mta* mutants via miR393b. **a)** Expression of auxin responsive reporter *proDR5:GUS* construct in *mta* mutants background is reduced upon treatment with 2,4 Dichlorophenoxyacetic acid (2,4D). Strong inhibition of auxin response after induction is seen in *mta* mutant. Scale bars (bottom right) = 2mm. **b)** miR393b level is downregulated in *mta* mutant (sRNA sequencing data) as compared to WT plants (left panel) (n=3), miR393b carries m^6^A mark (m^6^A-IP followed by RT-qPCR, middle panel) and pri-miR393b is bound by MTA (MTA-GFP RIP, right panel, n=3). Graphs represent fold change values between WT and *mta* mutant plants (1^st^ and 2^nd^ graph) and GFP and MTA-GFP plants (3^rd^ graph). **c)** *Nicotiana* leaves were agroinfiltrated with single constructs (MTA, catalytically inactive MTA (ΔMTA) and pri-miR393b; lane 2, 3 and 4 respectively) or a combination of two constructs (primiR393b +MTA (lane 5) and primiR393b + ΔMTA (lane 6)). Levels of miR393b were checked for each using northern blotting. Mock represents transfection with no plasmid control and U6 serves as loading control. Numbers above the last 3 lanes represent relative amounts of miR393b in presence of MTA (1.4, penultimate lane) and in presence of ΔMTA (2.3, last lane) as compared to pri-miR393b alone.

miR393b is also involved in auxin response regulation, and in mutants lacking miR393b, auxin signaling is reduced^46^. In our sRNA sequencing data, we found that miR393b is downregulated in *mta* mutant plants, although we could not detect its precursor in our m^6^A-IP Seq. However, because of its recognized importance in regulating auxin responses, we further investigated the requirement of m^6^A for miR393b formation. We confirmed that pri-miR393b is m^6^A methylated using m^6^A-IP followed by RT-qPCR. Immunoprecipitation of RNA bound to MTA also confirmed MTA’s binding to this precursor (**Fig. 5b)**. To further demonstrate that the lack of miR393b is infact caused by the lack of MTA, we designed a transient expression assay in *Nicotiana tobaccum*. We used constructs designed to express pri-miR393b, MTA or a catalytically inactive version of MTA, ΔMTA (D482A). ΔMTA was prepared by primer induced point mutation resulting in the following change: Aspartic acid at position 482 to Alanine (D482A) at the catalytic **D**PPW motif ^47^ leaving the protein catalytically inactive. These were introduced individually or in combination into *Nicotiana* leaves. We then assessed mature miR393b levels by northern blot and found that the pri-miRNA393b transgene produces ∼2.3 times more miR393b when it is present with MTA while this effect was abolished to a large extent when MTA was replaced by the catalytically inactive version (ΔMTA) (**Fig. 5c**). Thus, we show that the reduced auxin response in *mta* mutants may be partially caused by the regulatory defects of miR393b biogenesis caused by lack of MTA.

## Discussion

Here we provide the first evidence that, as well as influencing mRNA metabolism, m^6^A methylation is present in pri-miRNAs and affects miRNA biogenesis in *Arabidopsis.* We show that lower levels of MTA (and hence m^6^A) lead to a global reduction in miRNA levels whereas the pri-miRNAs tend to accumulate. This anti-correlation of pri-miRNAs with miRNAs is similar to what is seen in *Arabidopsis* mutants of genes encoding proteins involved in early stages of miRNA biogenesis. In contrast, mutants of HEN1 (a protein that is involved in methylation of miRNA/miRNA* duplex at the 3’ ends^48^, thus acting at later stages of miRNA biogenesis), do not show accumulation of pri-miRNAs whereas downregulation of miRNAs can be seen in *hen1* mutant plants^39,49^.

Since MTA is a known mRNA methyltransferase and m^6^A is abundant in mRNA, we considered the possibility that the observed affects were indirect and a result of altered metabolism of other miRNA biogenesis proteins. We used two different and independent approaches to exclude this possibility. MTA-GFP mediated RNA immunoprecipitation and m^6^A-IP experiments revealed that at least for a set of miRNAs, this effect is direct and is a result of MTA binding pri-miRNAs and introducing m^6^A marks.

While a direct effect of m^6^A on miRNA biogenesis is clear, we could not overlook the possibility that MTA could also affect miRNA biogenesis by interacting with some other miRNA biogenesis related proteins. This assumption is supported by the fact that MTA interacts with RNA Pol II as shown by our PLA results. MTA interacts with RNA Pol II phosphorylated at Ser5 and Ser2. This suggest that MTA is associated with the RNA Pol II from the initiation of transcription and is present during the elongation period, modifying growing transcripts (review ^50^). Furthermore, we show that MTA interacts with TGH, which indeed points to MTA being involved in early stages of miRNA biogenesis. Upon comparison of our data with previously published data regarding miRNA levels in *tgh* mutant we found a 45% overlap within downregulated miRNAs in *tgh* and *mta* mutants, despite data being from two different tissue sets^41^. These common miRNAs could be targets of m^6^A methylation facilitated by the MTA-TGH interaction; however, the exact mechanism of this interaction-related alteration of miRNA biogenesis requires further investigation. As TGH is a major miRNA biogenesis related protein, its effects on miRNA levels are more severe than those observed in relation to MTA indicating that MTA might affect biogenesis of a specific set of miRNAs whose selection might be assisted by TGH.

Finally, the reduced auxin responsiveness of m6A writer mutants may be partially explained by altered miR393b levels. We show that pri-miRNA393b is m^6^A methylated by MTA and levels of miR393b are lower in *mta* mutant. miR393b is involved in homeostasis of *AUX/IAA* genes that are also regulated by a complex feedback loop. miR393b has been shown to accumulate in leaves and is induced in response to auxin. Expression of the auxin responsive *DR5pro:GUS* reporter is reduced in *miR393b* mutant plants and a similar reduction is seen with the same reporter in the *mta* background^46,51,52^.

Our findings show that apart from methylating mRNAs, MTA is also a player in miRNA biogenesis. Similar to animal METTL3, MTA methylates pri-miRNAs, presumably at early stages of transcription and in its absence pri-miRNAs are accumulated and miRNA levels are downregulated. This altered miRNA biogenesis, including that of miR393b, may contribute to the reduced auxin response in m^6^A writer mutants.

## Methods

### Plant material

*Arabidopsis thaliana* WT Col-0, *mta* mutants^28^, MTA-GFP (*p35S:MTA-GFP* obtained from Dr. Rupert Fray (University of Nottingham)) and GFP plants were sown on Jiffypots® and stratified for 2 days in dark at 4°C. Thereafter, the plants were grown in plant growth chambers at 22°C with 16h light and 8h dark cycles (50–60% humidity, 150–200 µmolm^-2^ s^-1^ photon flux density). Rosette leaves from 4 weeks old plants were harvested, immediately flash frozen in liquid nitrogen and used, or stored at −80°C until further use.

### Small RNA sequencing and analysis

Total RNA from 4 weeks old *Arabidopsis thaliana* WT Col-0 and *mta* mutant plants was isolated using Direct-zol™ RNA kit (Zymo Research). RNA was quantified by Qubit RNA Assay Kit (Life Technologies) and quality by gel electrophoresis on Agilent Bioanalyzer 2100 system. 10 µg of RNA was separated on denaturing 15% PAA gel and small RNA fractions were cut out and purified from the gel. Libraries were prepared with TruSeq Small RNA Library Preparation Kit (Illumina) as per manufacturer’s instructions and sequencing was performed at Fasteris, Geneva, Switzerland on HiSeq 4000platform. The adapter Sequence was removed from the obtained raw reads with FASTX-Toolkit. Clean reads were mapped to all mature *A. thaliana* miRNAs from miRBase (release 22)^53^ using script: countreads_mirna.pl^54^. The script was applied to each fastq file for every biological replicate (n = 3). Subsequently, table with raw read counts was used to calculate fold change and false-discovery rate (FDR) with the usage of R-package NOISeq^55^. Normalization was included in the pipeline of NOISeq analysis.

### m^6^A-RNA immunoprecipitation of pri-miRNAs and sequencing

Total RNA from 4 week old *Arabidopsis* WT Col-0 and *mta*mutants^28^ was extracted as described above followed by polyA enrichment using NucleoTrap® mRNA isolation kit (Macherey-Nagel). Immunoprecipitation was performed with EpiMark N6-Methyladenosine Enrichment Kit (NEB) as per manufacturer’s instructions. Two RNA controls from the kit were spiked into RNA samples before immunoprecipitation (after polyA enrichment). Libraries from input and immunopreciptated RNA were prepared with SENSE Total RNA-Seq Library Prep Kit (Lexogen) and pair-end sequencing was performed at Fasteris, Geneva, Switzerland on HiSeq4000 platform. Raw reads were trimmed (first 30 nucleotides) with FASTX-Toolkit, adapters were removed using Trimmomatic^56^ andrRNA Sequences where removed with Bowtie^57^. Clean reads were aligned to the *Arabidopis* TAIR10 reference genome using HISAT2^58^ and the overall alignment rate was 89-92% for each sample. Transcript-level abundance for RNA in clean reads was estimated using Salmon^59^. Multifasta file with *Arabidopsis thaliana* reference transcripts (cDNA and ncRNA) for quasi-mapping was additionally supplemented with pri-miRNA transcirpts described in mirEX^2^ database^33^ and with m^6^A controls spike-in Sequences. TPM values in input and IP samples where normalized with the use of unmodified RNA spike-in control. Enrichment was calculated using TPM value with the following formula (wt IP/input) / (mta IP/input).

### Data submission and availability

The data obtained in this study (small RNA and m6A–IP Seq) has been deposited under NCBI GEO accession GSE122528.

### RNA immunoprecipitation (RIP)

Transgenic *Arabidopsis* line with *p35S:MTA-GFP* was grown along with GFP *Arabidopsis* line. RIPwas performed as described by Raczynska*et. al.*^60^, briefly, 4 week old leaves were crosslinked by vacuum infiltration in 1% formaldehyde for 10 min, quenched with 125mM glycine and frozen in liquid nitrogen. Nuclear fraction was isolated from this plant material following the protocol described by Bowler *et. al.*^61^and Kaufmann *et. al.*^62^. Isolated nuclei were sonicated at 4°C using Bioruptor® (Diagenode) at high intensity for 2 cycles (30 sec ON/30 sec OFF). GFP-Trap®_MA (Chromotek) magnetic beads were used to immunoprecipitate MTA-GFP or GFP fractions and RNA was isolated from input and IP samples. After DNAse treatment of RNA, it was primed with oligo(dT)primers for cDNA production and pri-miRNA levels were analyzed using SYBR® Green (Thermo) based qRT-PCR. 3 replicates each of MTA-GFP and GFP plants were analyzed and statistical significance was calculates using students T-test. Negative control (*AT2G40000)* was selected from data obtained from m^6^A-IP in a way that the gene transcript was not enriched in WT-IP after immunoprecipitation as compared to WT-Input, meaning that the transcript is likely not m^6^A methylated.

### Immunolocalization

The immunolabeling experiments and Duolink PLA fluorescence protocol were performed on isolated nuclei of 4-weeks old *Arabidopsis thaliana* leaves (WT, plants with MTA-GFP expression or GFP expression). Before isolation, the leaves were fixed in 4% paraformaldehyde in phosphate-buffered saline (PBS), pH 7.2, for 1 hour, washed 3 times in PBS in RT and then performed nuclei isolation protocol according to the method of Pontvianne *et al.*^63^

### Double Immunodetection of RNA Pol II and MTA

Prior to the assay, the cells were treated with PBS buffer containing 0,1% Triton X-100 for cell membrane permeabilization. MTA was localized by applying primary rabbit antibodies targeting GFP (Agrisera), diluted 1:200. For the localization of total fraction of RNAPII we used mouse 8WG16 antibody (anti-RNA polymerase II CTD repeat YSPTSPS, Abcam, diluted 1:200), according to protocols provided by Kołowerzo-Lubnau *et al*^64^. For RNA Pol II phosphorylated at serine 5 in CTD domain rat 3E8 antibody was used (Chromotek), diluted 1:100. For RNA Pol II phosphorylated at serine 2 in CTD domain rat 3E10 antibody was used (Chromotek), diluted 1:100. Primary antibody incubations (in 0,01% acetylated BSA in PBS) were performed in a humidified chamber overnight at 11°C. After washing with PBS, the slides were incubated with secondary goat anti-mouse antibodies: Alexa Fluor plus 488 (Thermo Fisher, diluted 1:200), or goat anti-rat antibodies Alexa Fluor 488 (Thermo Fisher, diluted 1:200) and goat anti-rabbit antibodies Alexa Fluor plus 555 (Thermo Fisher, diluted 1:200). The secondary antibodies were diluted in PBS containing 0,01% acetylated BSA and incubated at 37 °C in a humidified chamber for 1h. After the double labeling assay, the slides were stained for DNA detection with Hoechst 33342 (Life Technology, USA) and mounted in ProLong Gold antifade reagent (Life Technologies).

### Statistical analysis

Correlation analysis was performed with the use of Pearson’s correlation coefficient, Spearman’s rank correlation and ICQ value by using a free software ImageJ from the National Institute of Health in USA and its plugin coloc2 which is the analysis options of the expanded ImageJ version Fiji.

### Proximity Ligation Assay (PLA)

As for the immunofluorescence method, prior to the PLA assay, the cells were treated with PBS buffer containing 0,1% Triton X-100 for cell membrane permeabilization. During the PLA method, a 3-stage protocol was used using AB290 primary rabbit antibodies to GFP (Abcam, dilution 1: 100) and rat antibodies to RNA Pol II with serine 2/5 modification (Chromotech, dilution 1: 200). SubSequently, the Alexa Fluor 488 (Thermo Fisher, diluted 1:200) secondary goat anti-rat was used. Incubation with primary and secondary antibodies was carried out in a solution of 0.1% acBSA in PBS buffer in a humidified chamber at 4°C. Then, the PLA protocol was used. In situ PLA detection was carried out using the appropriate DUOLINK in situ Orange Kit Goat/Rabbit obtained from Sigma-Aldrich (Saint Luis, USA) according to the protocol of the manufacturer. After IF step cells were, washed with PBS, and then subjected to blocking using the DUOLINK blocking solution at 37°C in a humidified chamber for 60 min. After two washes with wash buffer A for 5 min each, secondary/tertiary antibodies (DUOLINK anti-rabbit PLA-plus probe, DUOLINK anti-goat PLA-minus probe) were added and incubated at 37°C for 1 h (at a dilution of 1:40 in 40 µl DUOLINK antibody diluents). The slides were washed two times with wash buffer A for 5 min each, then followed by addition of the ligation mix and incubation at 37°C for 30 min, followed by another two washes with Wash buffer A. Finally, the amplification reaction was carried out at 37°C for 100 min. Subsequently, the slides were washed twice with Washbuffer B (10 min. each), and once with 0.01×Washbuffer B 1 min. After the PLA assay, the slides were stained for DNA detection with Hoechst 33342 (Life Technology) and mounted in ProLong Gold antifade reagent (Life Technologies). This type of 3-step reactions allowed to locate both the total pool of RNAPol II ser2 / 5 (green fluorescence) as well as those that are associated with MTA (orange spots of fluorescence). Control reactions were performed on isolated nuclei obtained from wild-type plants and those expressing only GFP proteins.

### Microscopy

The results were registered with Leica SP8 confocal microscope using lasers emitting light at wavelengths of 405, 48 and 561 nm. For Leica confocal microscope an optimized pinhole, long exposure time (200 kHz) and 63X (numerical aperture, 1.4) Plan Apochromat DIC H oil immersion lens was used. Images were collected sequentially in the blue Hoechst 33342), in the green (Alexa 488 fluorescence) and red (Alexa 555, PLA Orange) channels. To minimize bleed-through between fluorescence channels, we employed low laser power (0.4–5% of maximum power) and single-channel collection. For bleed-through analysis and control experiments, Leica SP8 software was used.

### Yeast two hybrid assay

cDNA clones of MTA, TGH, SE, CBP20, CBP80, HYL1 and DDL1 were prepared in pGADT7 and pGBKT7 and the experiment was performed following the protocol of Matchmaker® Gold Yeast Two-Hybrid system (Clontech). Briefly, *Saccharomyces cerevisiae* strain Y2HGold was co-transformed with 700 ng of each plasmid encoding fusion proteins according to the manufacturer’s protocol, and spread on three different selective media: Double dropout medium (SD/-Leu/-Trp), Quadruple dropout medium (SD/-Leu-Trp-His-Ade) and QDO+ (QDO + X-α-Gal + Aureobasidin A) After growing of yeast in selective media at 28°C for 3 days interactions were identified by bluish green colonies on QDO+ medium (n = 3).

### Microscopy and FRET-FLIM

Coding sequences of MTA and TGH were first cloned into pENTR™/D-TOPO® vectors (Thermo Fischer). Gateway LR Clonase II Enzyme Mix (ThermoFisher Scientific) was used to clone these sequences into destination vectors with GFP/RFP tag and *UBQ10* promoter. Protoplasts were isolated from 3-4 weeks old WT *Arabidopsis* leaves using Tape-*Arabidopsis* method that were then processed as described in ^65^. Prepared protoplasts were then transfected with the expression constructs of tagged proteins using a PEG/Ca^2+^ solution as described in ^65^. After 7 hours of incubation, cells were visualized under confocal microscope (Nikon A1Rsi) and FRET-FLIM measurements were undertaken using PicoHarp300-Dual Channel SPAD system (PicoQuant) in combination with Nikon A1Rsi microscope. 9 cells were analyzed for each sample and statistical significance was calculated using student’s t-test.

### GUS staining for auxin response analysis

14 days old seedlings of WT and *mta* mutants with*DR5pro:GUS* reporter gene construct were incubated in ½ MS medium containing 10 µM 2,4-D in ethanol or ethanol alone as a control for 8h. After GUS staining the pictures were taken under Leica M60 Stereo Microscopy.

### pri-miR393b, MTA and ΔMTA constructs

pri-miR393b (Sequence from mirEX^2^, http://www.combio.pl/mirex2) and MTA cDNA (*AT4G10760*) were amplified and cloned into pENTR™/D-TOPO® vectors (Thermo Fischer). ΔMTA was prepared by primer induced point mutation resulting in the following change: Aspartic acid at position 482 to Alanine (D482A) at the catalytic **D**PPW motif. For transient expression in *Nicotiana* these sequences were then cloned into pMDC32 vector using Gateway® (Thermo) cloning system.

### *Nicotiana tobaccum* transient expression and northern blotting

pri-miR393b, MTA and ΔMTA constructs were transformed into *Agrobacterium tumifaciens* using electroporation. After verification of constructs in *Agrobacterium* by sequencing, *Nicotiana* leaves were transformed as described by Bielewicz *et. al.*^66^. Leaves were transformed either with pri-miR393b, MTA and ΔMTA alone or in pairs (pri-miR393b + MTA, pri-miR393b + ΔMTA); leaves were harvested after 72h and RNA isolation (as described above) was followed by northern blotting.

### Northern Blotting

Northern blotting was performed as described in^67^, briefly: 30 µg of RNA (per sample) isolated from transfected *Nicotiana* leaves was loaded on 8M denaturing urea gel (Polyacrylamide 15%) in TBE buffer. RNA was then transferred onto Amersham Hybond-NX nitrocellulose membrane (GE Healthcare) using a Trans-Blot Electrophoretic Transfer Cell (Bio-Rad) and fixed using CL-1000 Ultraviolet Crosslinker (UVP). Pre-hybridization and hybridization were performed in hybridization buffer (3.5% SDS, 0.375 M sodium phosphate dibasic, 0.125 M sodium phosphate monobasic) at 42°C with DNA oligo probes (Sigma) labeled with γ^32^P ATP. U6 was used as a loading control. After washing, the blots were exposed for upto 3 days to phosphorimaging screen (Fujifilm) and results were visualized with Fujifilm FLA5100 reader (Fujifilm) and quantified using Multi Gauge V2.2 (Fujifilm).

## Acknowledgments

This work was funded by KNOW Poznan RNA Centre (grant no. 01/KNOW2/2014) and Polish National Science Centre (grants no. UMO-2017/27/N/NZ1/00202, UMO-2016/23/B/NZ9/00862, UMO-2013/10/A/NZ1/00557). Work in R.F.’s lab was supported by the Biotechnology and Biological Sciences Research Council (grant BB/M008606/1).

## Author contribution

S.S.B participated in experimental design, performed the majority of experiments and participated in writing the MS; D.B participated in experimental design, performed bioinformatical analysis and participated in discussion, analysis of results and MS writing; N.G contributed towards tobacco transient expression assays; T.G performed immunolocalization assays; D.S performed PLA assays and participated in writing MS; Z.B participated in m6A IP; R.G.F participated in m6A IP and auxin response experiment; M.B, J.D and L.S participated in experiments and discussion; A.J participated in experimental design, discussion and analysis of results and writing the MS; Z.S.K conceived the idea and designed experiments, participated in discussion and results analysis as well as writing the MS

