## Supplementary for "mRNA adenosine methylase (MTA) deposits m^6^A on pri-miRNAs to modulate miRNA biogenesis in *Arabidopsis thaliana*"

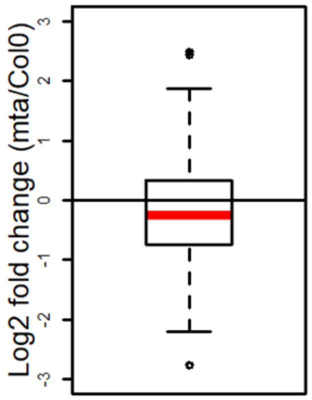


**Figure S1 miRNA biogenesis is impaired in *mta* mutants.** An overall downregulation of miRNAs in *mta* mutants (as compared to WT plants) as determined by sRNA sequencing is visualized by a box plot. Red bar represents median value.

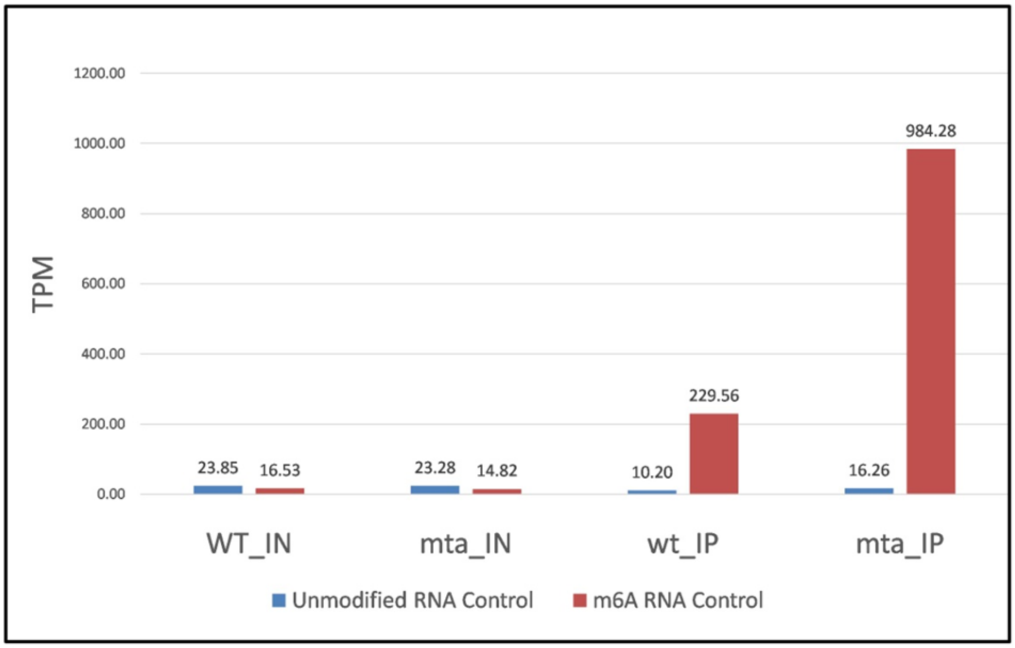


**Figure S2 RNA spike containing m$^6$A modification is efficiently recognized by m6a antibodies.** Enrichment of m$^6$A modified RNA control (red colour) that was spiked in the samples can be seen in m$^6$A immunoprecipitated samples (WT_IP and *mta*_IP). Unmodified RNA control (blue colour) is not enriched in m$^6$A immunoprecipitated samples (WT_IP and *mta*_IP). WT_IN and *mta*_IN represents input samples whereas WT_IP and *mta*_IP represent immunoprecipitated samples. Numbers above the columns represent transcripts per million of each sample respectively.

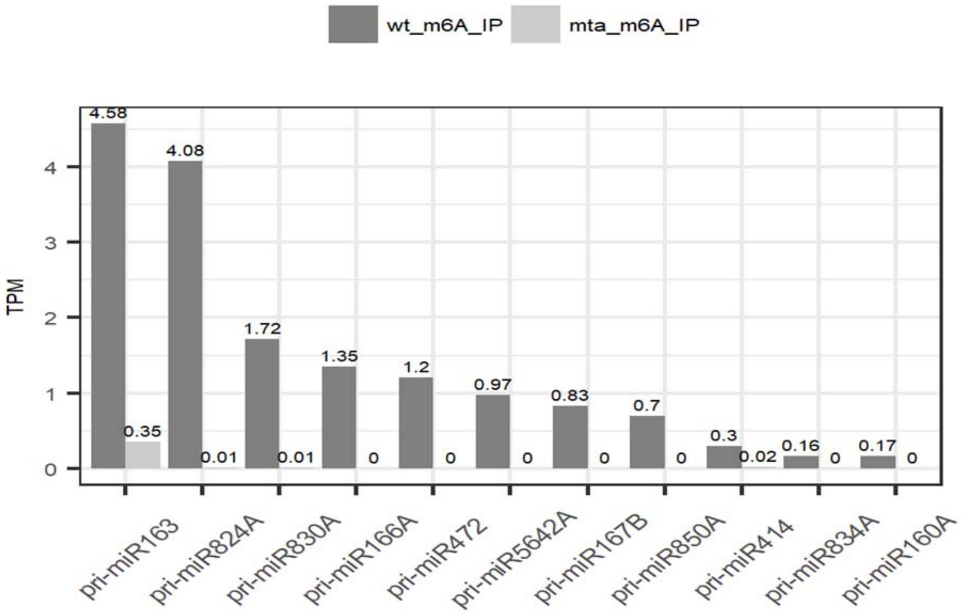


**Figure S3 pri-miRNA's are m$^6$A methylated:** Enrichment of *MIR* gene precursors identified by m$^6$A IP Seq that can be seen in immunoprecipitated samples from WT plants (wt_m$^6$A_IP) but not from *mta* mutants (*mta*_m$^6$A_IP). Numbers represent transcripts per million (TPM).

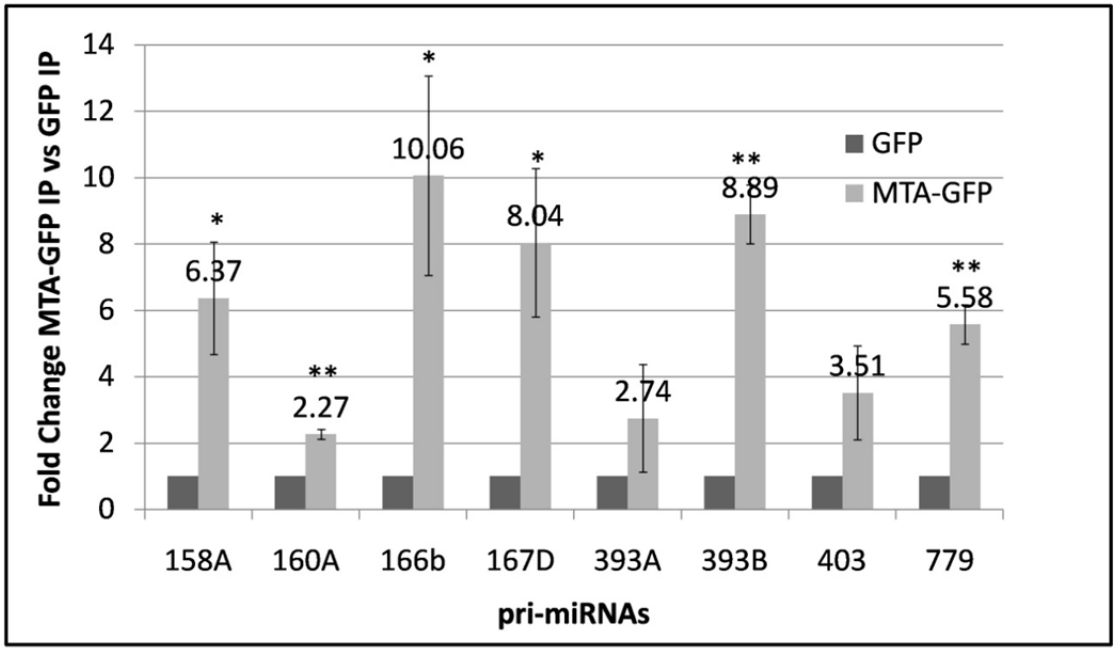


**Figure S4 Enrichment of pri-miRNAs in MTA-GFP IP samples.** 8 randomly selected pri-miRNAs show enrichment in MTA-GFP samples after RIP experiment. Values represent fold change and * = p-value < 0.05, ** = p-value < 0.005. Error bars represent standard deviation (n=3).

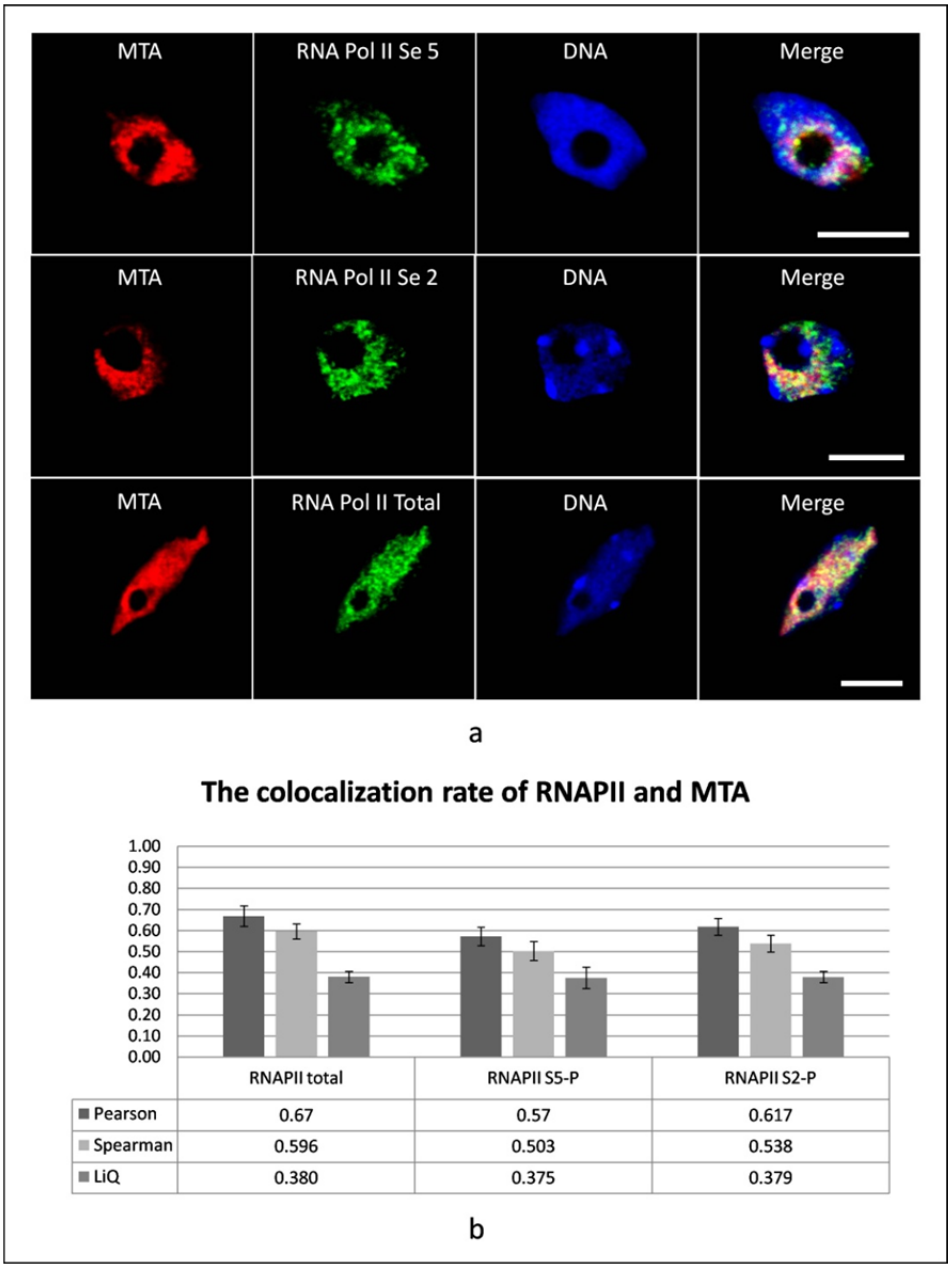


**Figure S5 MTA co-localizes with RNA Pol II.** Immunolocalization images show co-localization of MTA with RNA Pol II phosphorylated at Serine 5 (RNA Pol II Se 5), Serine 2 (RNA Pol II se 2) and total RNA Pol II (RNA Pol II). MTA is detected by Alexa Fluor plus 555, RNA Pol II by Alexa Fluor plus 488 and DNA is stained by HOECHST. Merge column represents merge of all three columns. Scale bars = 5µm **b)** Colocalization rate of MTA and RNA as measured using Pearson, Spearmans and LiQ coefficients. Bars represent various colocalization parameter scores and error bars depict standard deviation.

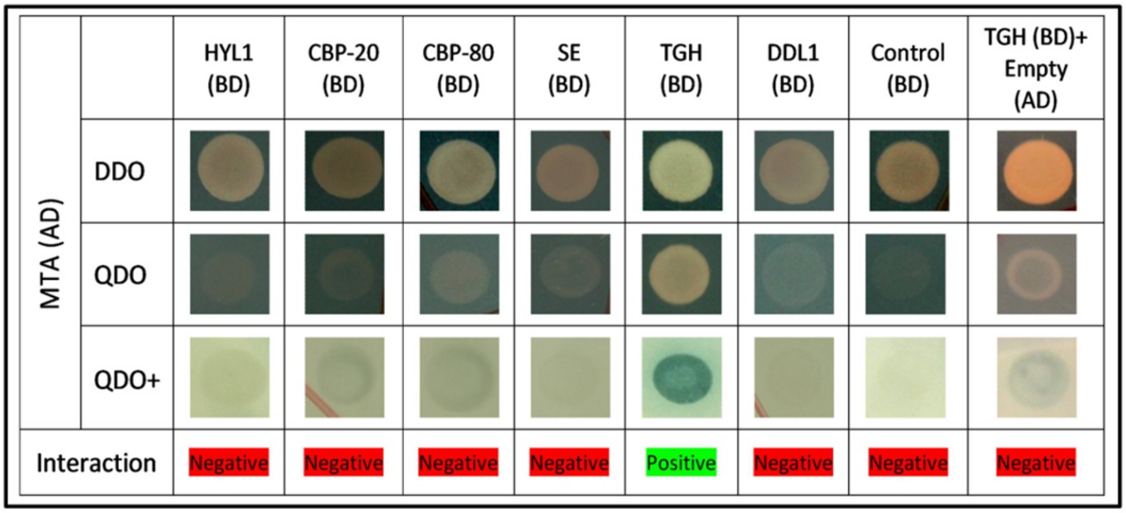


**Figure S6 MTA interacts with TGH.** Yeast two hybrid experiment between MTA fused with activation domain (AD) and selected miRNA biogenesis proteins fused to binding domain (BD) showed positive interaction between MTA and TGH. Bluish green colonies in quadruple dropout medium (QDO+) show positive interactions.

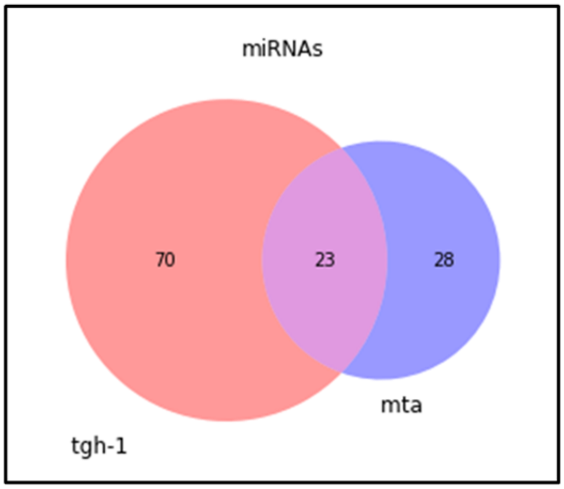


**Figure S7 Comparison of sRNA sequencing data from *mta* (rosette leaves) and *tgh-1***

**(inflorescence tissue) mutants.** 23 out of 51 miRNAs downregulated in *mta* mutant are common between *tgh* and *mta* mutant. Numbers represent miRNAs downregulated in these mutants.
